# Mutations in transcription factors that confer fluconazole resistance also confer reduced susceptibility to manogepix in *Candida auris*, *Candida albicans*, *Candida parapsilosis*, and *Candida glabrata* (*Nakaseomyces glabratus*)

**DOI:** 10.1101/2025.05.07.652603

**Authors:** Katherine S. Barker, Hoja P. Patterson, Joachim Morschhäuser, Christina A. Cuomo, Nathan P. Wiederhold, P. David Rogers

## Abstract

The fungal pathogen *Candida auris* is of global concern due to high levels of multidrug resistance and its propensity to cause infectious outbreaks. Over 90% of isolates are resistant to fluconazole, the most commonly prescribed antifungal world-wide. Fluconazole resistance is multifactorial with many isolates carrying mutations in the gene encoding the transcriptional regulator Tac1B, leading to increased expression of the gene encoding the ATP Binding Cassette (ABC) transporter Cdr1. Recently, a study examining *C. auris* in vitro resistance mechanisms to manogepix, a promising antifungal agent currently in clinical trials, found a *TAC1B* mutation that confers reduced manogepix and fluconazole susceptibility. We hypothesized that mutations in *C. auris TAC1B* and similar transcription factors in other *Candida* species that confer fluconazole resistance might also confer reduced susceptibility to manogepix. We measured manogepix susceptibilities for selected isolates and strains and found *C. auris TAC1B*, *C. albicans* and *C. parapsilosis TAC1*, and *C. glabrata PDR1* confer reduced manogepix susceptibility in a manner dependent on ABC transporters similar to Cdr1. Our findings raise the possibility of fluconazole and manogepix cross resistance for clinical isolates harboring mutations in these genes.

## Introduction

Invasive candidiasis is associated with significant morbidity and mortality. It is estimated that over 400,000 people die annually from invasive candidiasis (IC) with an average mortality rate of 35% (1). The majority of infections are caused by five *Candida* species: *Candida albicans*, *Candida glabrata* (*Nakaseomyces glabratus*), *Candida tropicalis*, *Candida parapsilosis*, and *Candida krusei* (*Pichia kudriavzevil*) (2). Moreover, *Candida auris* (*Candidozyma auris*) has emerged as a global health threat owing to its propensity to cause outbreaks in the healthcare environment and high rates of antifungal resistance (3, 4). Three classes of antifungal agents are useful for the treatment of IC; the echinocandins such as micafungin, the triazoles such as fluconazole, and the polyene amphotericin B. While the echinocandins have emerged as front-line therapy for the treatment of serious infectious due to *Candida* species, fluconazole is still widely used and remains the most commonly prescribed antifungal in the U.S. (5).

Fosmanogepix is a prodrug of the active moiety manogepix, which acts by inhibiting Gwt1, a key enzyme in the glycosylphosphatidylinositol pathway and biosynthesis of glycosylphosphatidylinositol (GPI) anchors, resulting in compromised cell wall integrity (6). With the exception of the less common species *C. krusei, C. inconspicua* and *C. kefyr*, manogepix exhibits broad spectrum fungistatic activity against *Candida* species (7). While formal susceptibility breakpoints have not been established, epidemiologic cutoff values have been suggested for manogepix as 0.004-0.008 µg/mL for *C. albicans*, 0.016 µg/mL for *C. parapsilosis* and *C. auris*, and 0.06 µg/mL for *C. glabrata* (8). In that study a correlation with fluconazole susceptibility was observed. One possible explanation is the contribution of efflux pumps as manogepix susceptibility has been shown to be influenced by *CDR11* and *SNQ2* encoding ATP binding cassette (ABC) transporters in *C. albicans* and *MDR1* encoding a Major Facilitator Superfamily (MFS) transporter in *C. parapsilosis* (9).

In *C. albicans*, a major driver of fluconazole resistance is overexpression of the ABC transporter gene *CDR1* due to activating mutations in the zinc cluster transcription factor *TAC1* (10). In *C. parapsilosis*, resistance is driven in part by overexpression *CDR1*, *CDR1B*, and *CDR1C*, due to activating mutations in *TAC1*, whereas in *C. auris* it is driven in part by overexpression of *CDR1* and to a lesser extent *MDR1*, due to mutations in *TAC1B* (11–13). In *C. glabrata*, fluconazole resistance is almost exclusively due to activating mutations in the zinc cluster transcription factor gene *PDR1* that lead to overexpression of the *CDR1*, *PDH1*, and *SNQ2* transporter genes alone or in combination (14–16).

Recently, in vitro evolution studies in *C. auris* identified a mutation in *TAC1B* in laboratory strains evolved to have reduced susceptibility to manogepix (17). This D865N amino acid substitution resulted in increased expression of *CDR1*, and loss of either *CDR1* or *TAC1B* resulted in increased manogepix susceptibility in these strains. These findings suggest that fluconazole-resistant *Candida* isolates with mutations in similar transcription factor genes may likewise affect manogepix susceptibility.

In order to assess the contribution of these genes to manogepix susceptibility and to determine the potential for cross resistance between fluconazole and manogepix, we determined the susceptibilities to manogepix in clinical isolates and strains from our collection that have mutations in *TAC1B* (*C. auris*), *TAC1* (*C. albicans* and *C. parapsilosis*) , and *PDR1* (*C. glabrata*) that drive overexpression of ABC drug transporters in these species and are known to contribute to fluconazole resistance.

## Results

### Mutations in *TAC1B* that confer increased resistance to fluconazole in *C. auris* confer reduced susceptibility to manogepix

To determine if mutations in *TAC1B* that confer increased resistance to fluconazole in *C. auris* influence susceptibility to manogepix, we first measured manogepix susceptibilities in the fluconazole-resistant clinical isolate Kw2999 which carries a mutation leading to the A640V substitution, its derivative strain 1c where the *TAC1B* gene was corrected to the wild-type sequence, and derivatives of 1c where mutations leading to A640V, A657V, and F862_N866del found in fluconazole-resistant clinical isolates were introduced (**Fig 1A**) (13). Isolate Kw2999 exhibited manogepix MICs of 0.03 µg/mL as compared to 0.008 µg/mL for strain 1c, a two-dilution difference in MIC. Introduction of these three mutations into strain 1c resulted in MICs of 0.03 µg/mL, 0.03 µg/mL, and 0.125 µg/mL representing two-, two-, and four-dilution differences in susceptibility. We then measured manogepix susceptibilities in another fluconazole-susceptible strain AR0387, its *TAC1B*^A640V^ derivative strain, fluconazole-resistant isolate AR0390 which harbors the *TAC1B*^A640V^ allele, and its wild-type *TAC1B* derivative strain (**Fig 1B**). Introduction of *TAC1B*^A640V^ into AR0387 reduced manogepix susceptibility by two dilutions from 0.008 µg/mL to 0.03 µg/mL whereas its correction to the wild-type sequence in AR0390 reduced manogepix susceptibility by two dilutions from 0.03 µg/mL to 0.008 µg/mL. All isolates and strains with a *TAC1B* mutation exhibited manogepix MICs at or above the proposed epidemiologic cutoff value (8). These data indicate that activating mutations in *TAC1B* that confer reduced susceptibility to fluconazole in *C. auris* clinical isolates likewise confer reduced susceptibility to manogepix.

**Figure 1.**
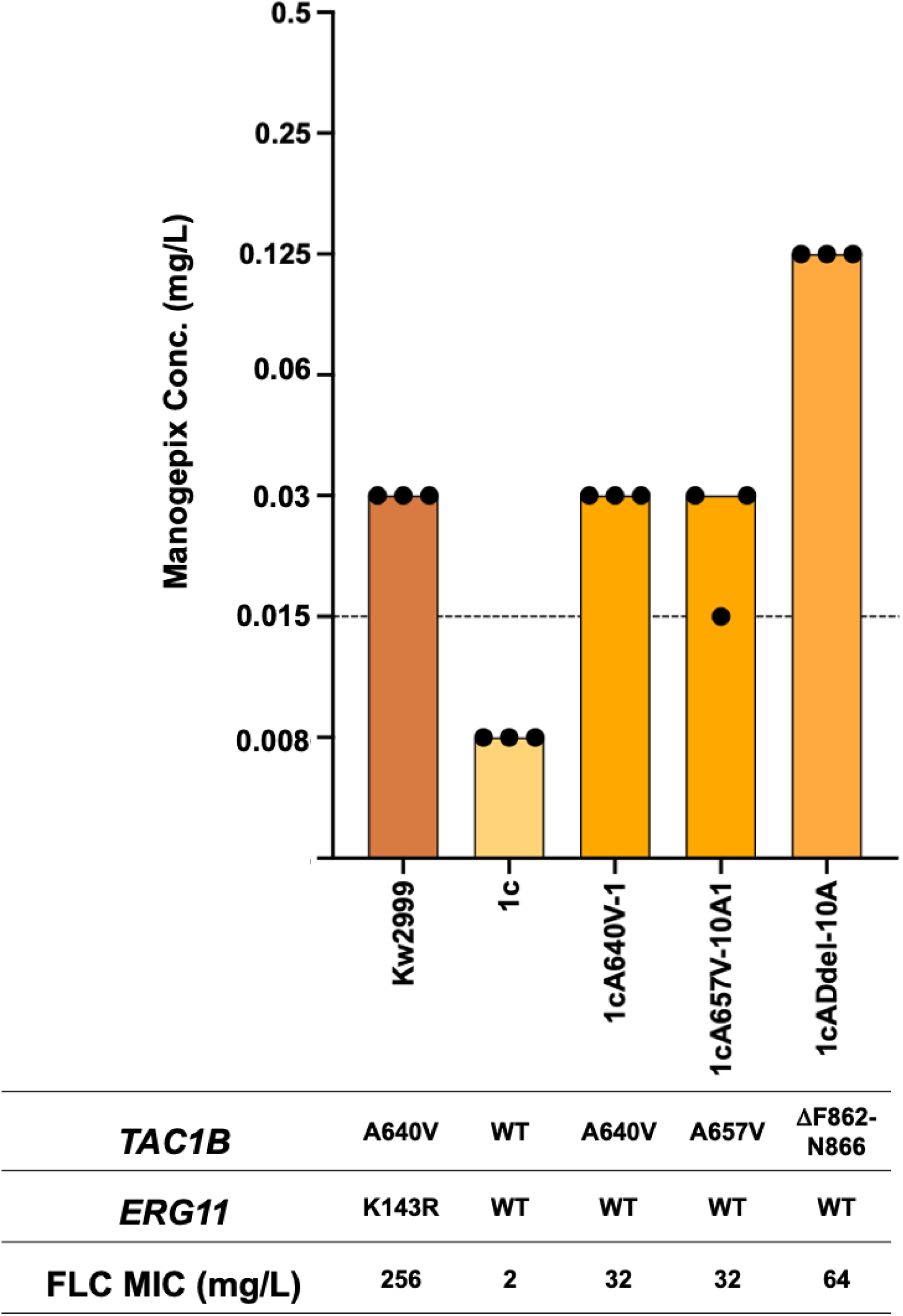
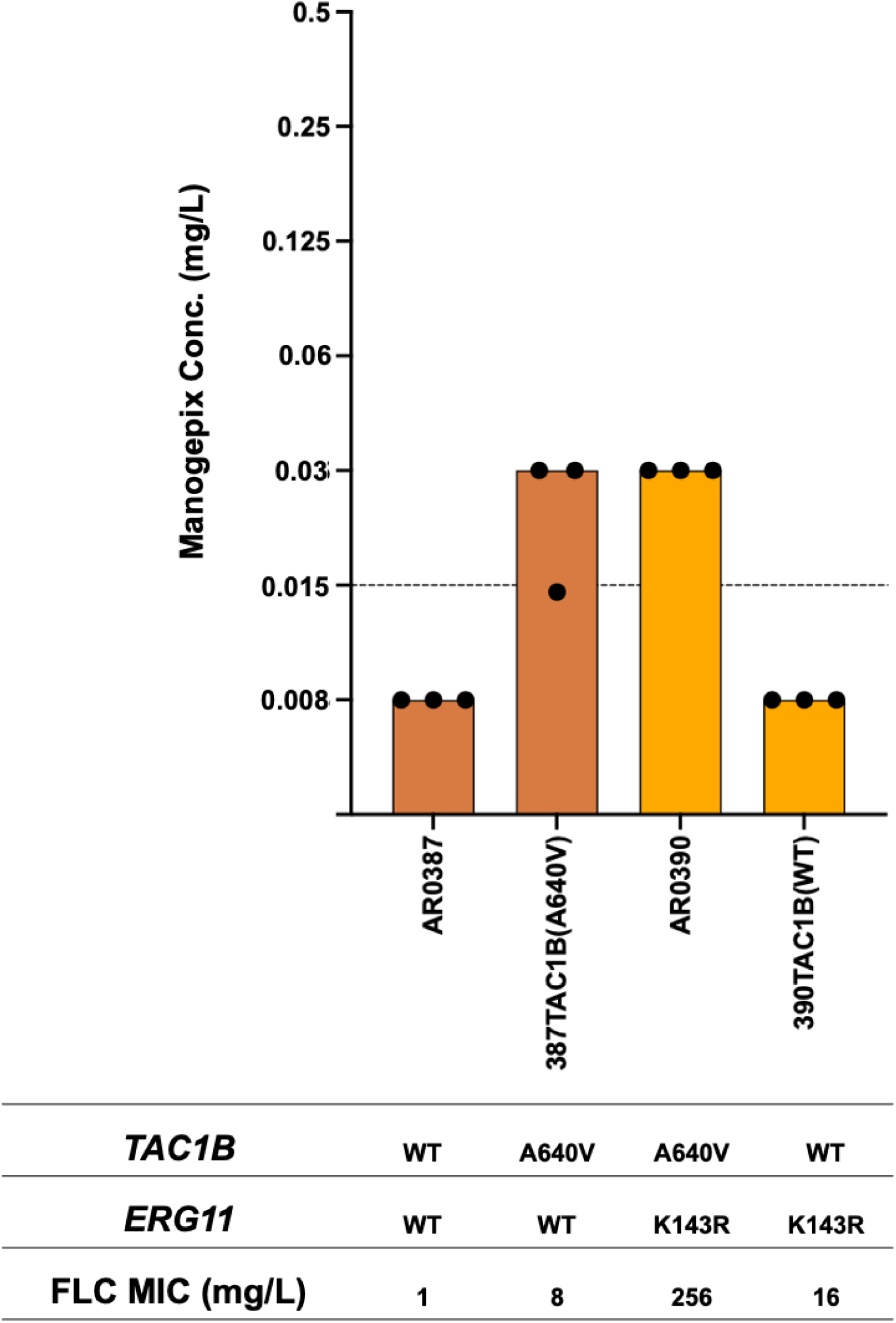
Manogepix MIC for (A) *Candida auris* clinical isolate Kw2999 and its derivative strains harboring either *TAC1B*^WT^ or most prevalent *TAC1B* mutant alleles, and (B) *C. auris* CDC AR Bank isolates AR0387 and AR0390 and their *TAC1B* allele-swapped derivative strains. Each bar in graph represents the modal MIC value, and each dot represents the measurements from three independent assays. Proposed epidemiologic cutoff value is indicated by the dashed line.

### *CDR1* but not *MDR1* drives reduced manogepix susceptibility in fluconazole resistant isolates of *C. auris* carrying mutations in *TAC1B*

In *C. auris*, *TAC1B*-mediated fluconazole resistance is driven by overexpression of both the *CDR1* and *MDR1* transporter genes (13). In isolates that are resistant to fluconazole due to mutations in *TAC1B*, disruption of *CDR1* confers increased fluconazole susceptibility. Disruption of *MDR1* alone has no effect on fluconazole susceptibility, but disruption of both *CDR1* and *MDR1* confers a greater increase in susceptibility than *CDR1* disruption alone. We therefore measured manogepix susceptibilities in a panel of strains carrying mutations in *TAC1B* where *CDR1* and *MDR1* have been disrupted (**Fig 2**). Disruption of *CDR1* in strains engineered to express *TAC1B* mutations leading to A640V, A657V, or F862_N866del resulted in a three-, three-, and five-dilution increase in manogepix susceptibility from 0.03 µg/mL, 0.03 µg/mL, and 0.125 µg/mL to 0.004 µg/mL for these strains respectively. Disruption of *MDR1* in these strains had no effect on manogepix susceptibility, and disruption of both *CDR1* and *MDR1* resulted in no greater increase in susceptibility than *CDR1* disruption alone. These data indicate that *CDR1* is the major driver of reduced manogepix susceptibility due to these *TAC1B* mutations.

**Figure 2.**
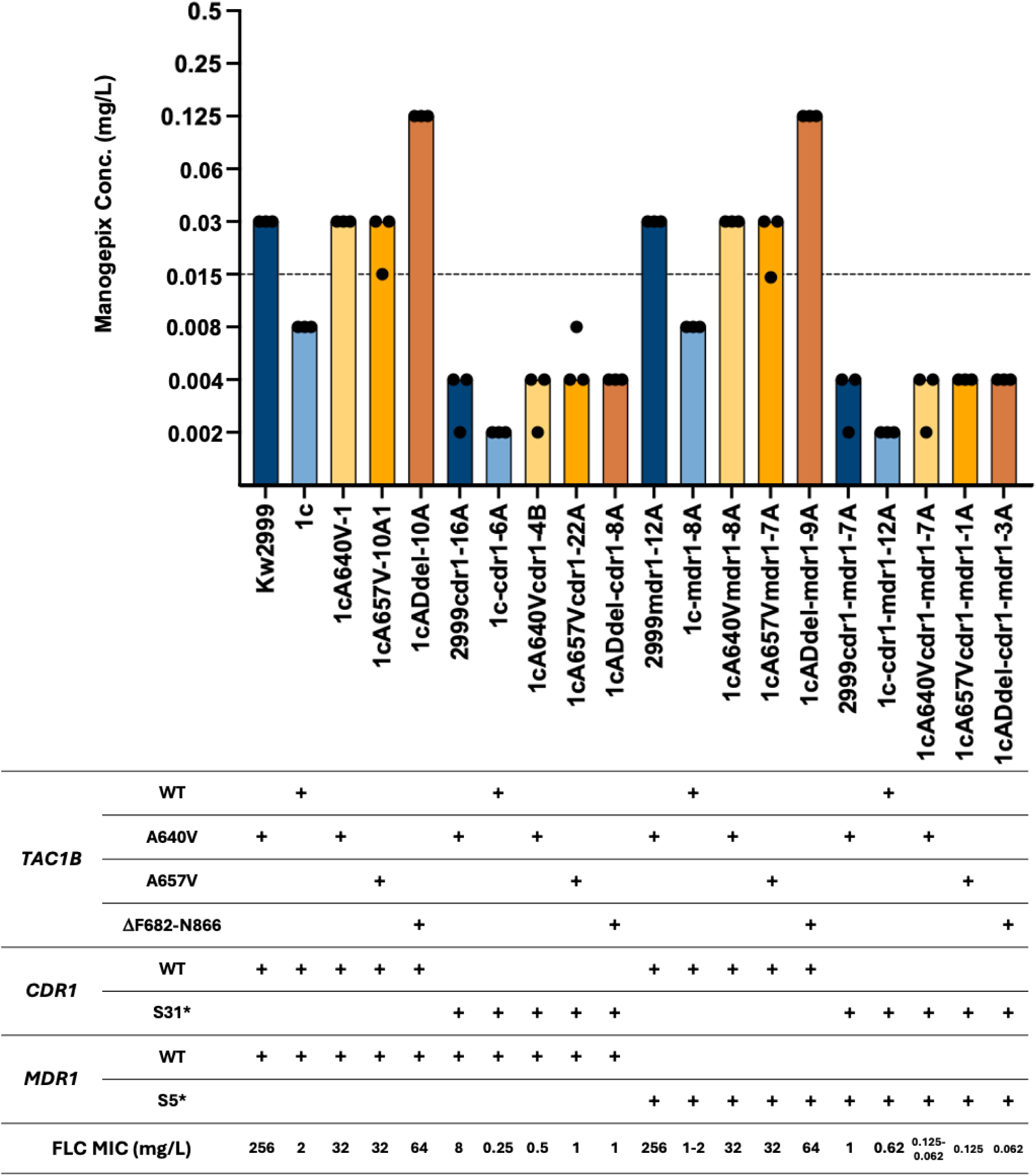
Manogepix MIC for *cdr1*-disrupted, *mdr1*-disrupted, or *cdr1*/*mdr1*-disrupted *TAC1B* mutant *C. auris* strains. Each bar in graph represents the modal MIC value, and each dot represents the measurements from three independent assays. Proposed epidemiologic cutoff value is indicated by the dashed line.

### Mutations in *TAC1* that confer increased resistance to fluconazole in *C. albicans* confer reduced susceptibility to manogepix

In order to determine if activating mutations in *TAC1* that contribute to fluconazole resistance in *C. albicans* also influence manogepix susceptibility, we determined manogepix susceptibilities in a panel of matched pairs of fluconazole-susceptible and - resistant clinical isolates where the resistant isolates carry activating mutations in *TAC1*. These isolates and strains have been shown previously to overexpress *CDR1* and *CDR2* (18–22). Each resistant isolate was obtained from the same patient as its matched susceptible counterpart over the course of fluconazole treatment. Fluconazole-resistant isolates Gu5, C56, TW17, and 5674 carry mutations leading to the G980E, N977D, A736V/Δ962-969, and N972D amino acid substitutions, respectively. Their respective manogepix MICs were 0.06 µg/mL, 0.06 µg/mL, 0.03 µg/mL, and 0.125 µg/mL which is a three-, two-, two-, and four-dilution increase in manogepix MIC, relative to their respective matched susceptible isolates (**Fig 3A**).

**Figure 3.**
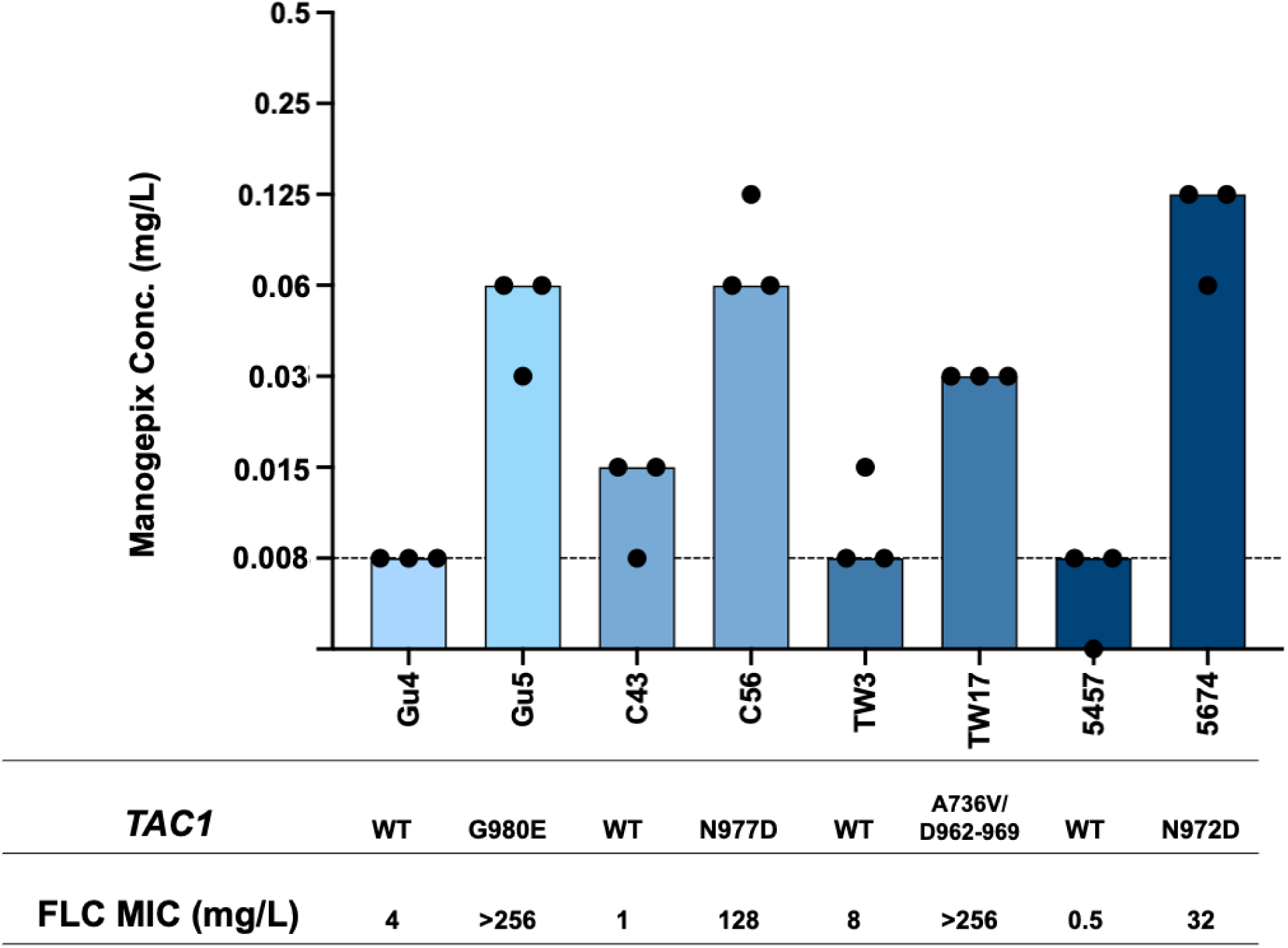
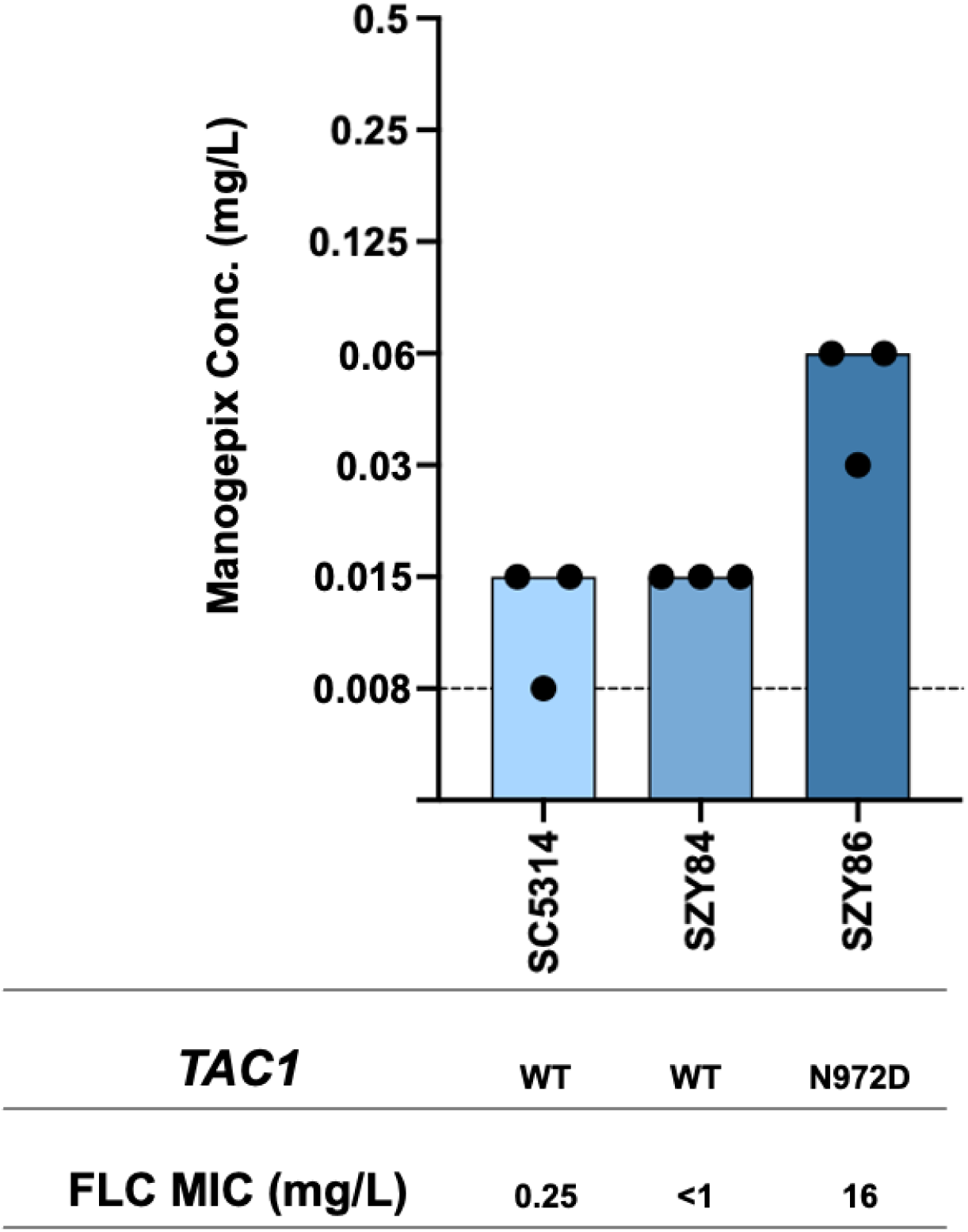
Manogepix MIC for (A) *C. albicans* matched set clinical isolates in which the fluconazole-resistant isolate harbors a *TAC1* mutation, and (B) *C. albicans* laboratory strain SC5314 and its derivative strains harboring a *TAC1* allele from fluconazole-susceptible isolate 5457 or a *TAC1* allele from fluconazole-resistant isolate 5674. Each bar in graph represents the modal MIC value, and each dot represents the measurements from three independent assays. Proposed epidemiologic cutoff value is indicated by the dashed line.

Strains SZY84 and SZY86 are derived from strain SC5314 where either a wild-type *TAC1*allele from fluconazole-susceptible clinical isolate 5457 or the *TAC1* allele with a mutation leading to the N972D substitution from resistant isolate 5674 was introduced into a *tac1*Δ/*tac1*Δ strain (22). SZY86 exhibited a 2-dilution increase in manogepix MIC to 0.06 µg/mL as compared to 0.015 µg/mL SZY84 (**Fig 3B**). These results indicate that activating mutations in *TAC1* that confer fluconazole resistance likewise result in reduced susceptibility to manogepix in *C. albicans*.

### A mutation in *TAC1* that confers increased resistance to fluconazole in *C. parapsilosis* confers reduced susceptibility to manogepix

In order to determine if mutations in *TAC1* in *C. parapsilosis* that confer fluconazole resistance influence susceptibility to manogepix, we measured manogepix susceptibilities in clinical isolate Cp35 which harbors a *TAC1*^G650E^ mutation as well as its derivative where its *TAC1* sequence was corrected to wild-type. Cp35 exhibited a MIC of 0.06 µg/mL, 2-dilutions higher than its *TAC1*^WT^ derivative (**Fig 4A**). This observation indicates that an activating mutation in *TAC1* that confers increased resistance to fluconazole in *C. parapsilosis* likewise confers reduced susceptibility to manogepix.

**Figure 4.**
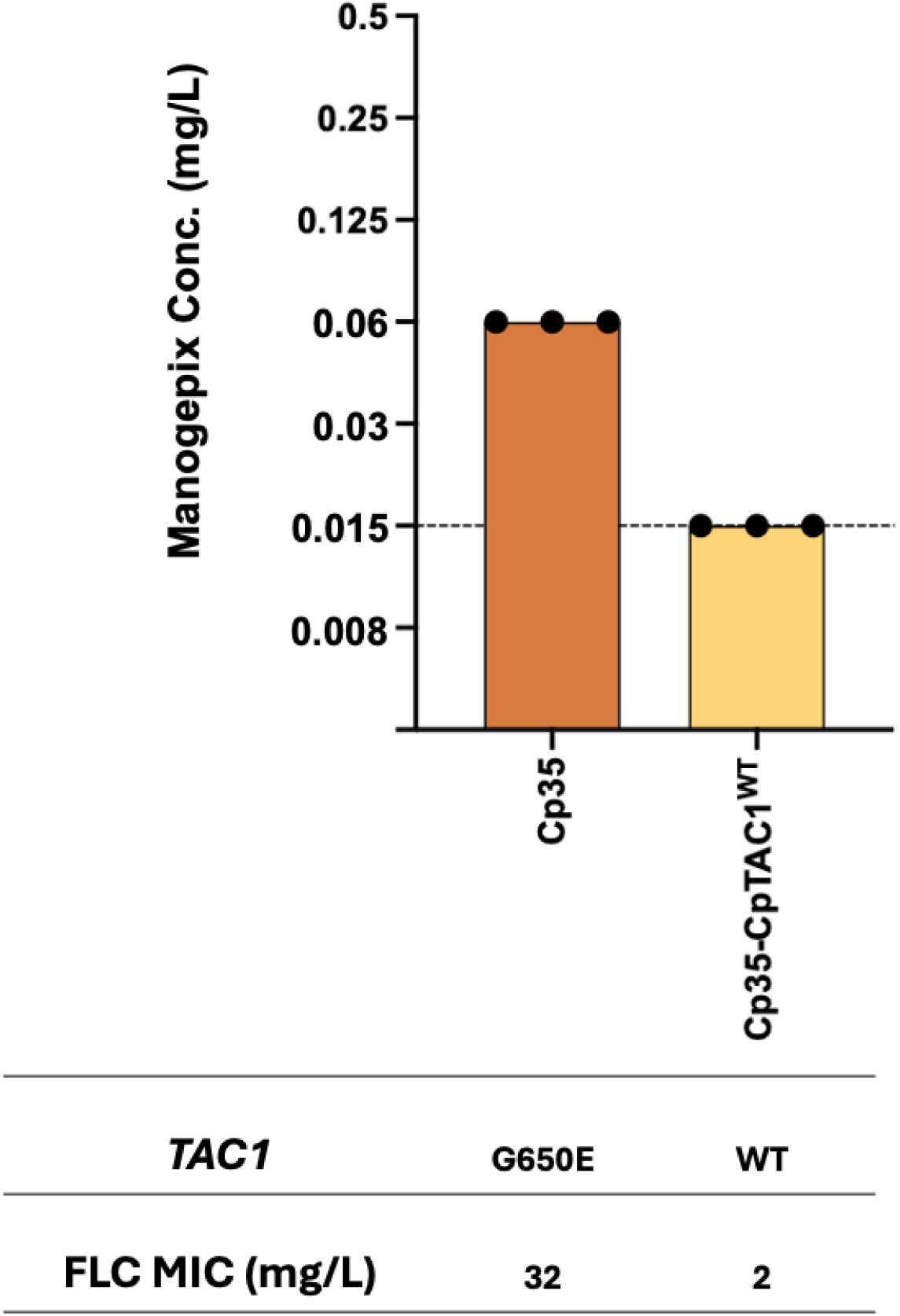
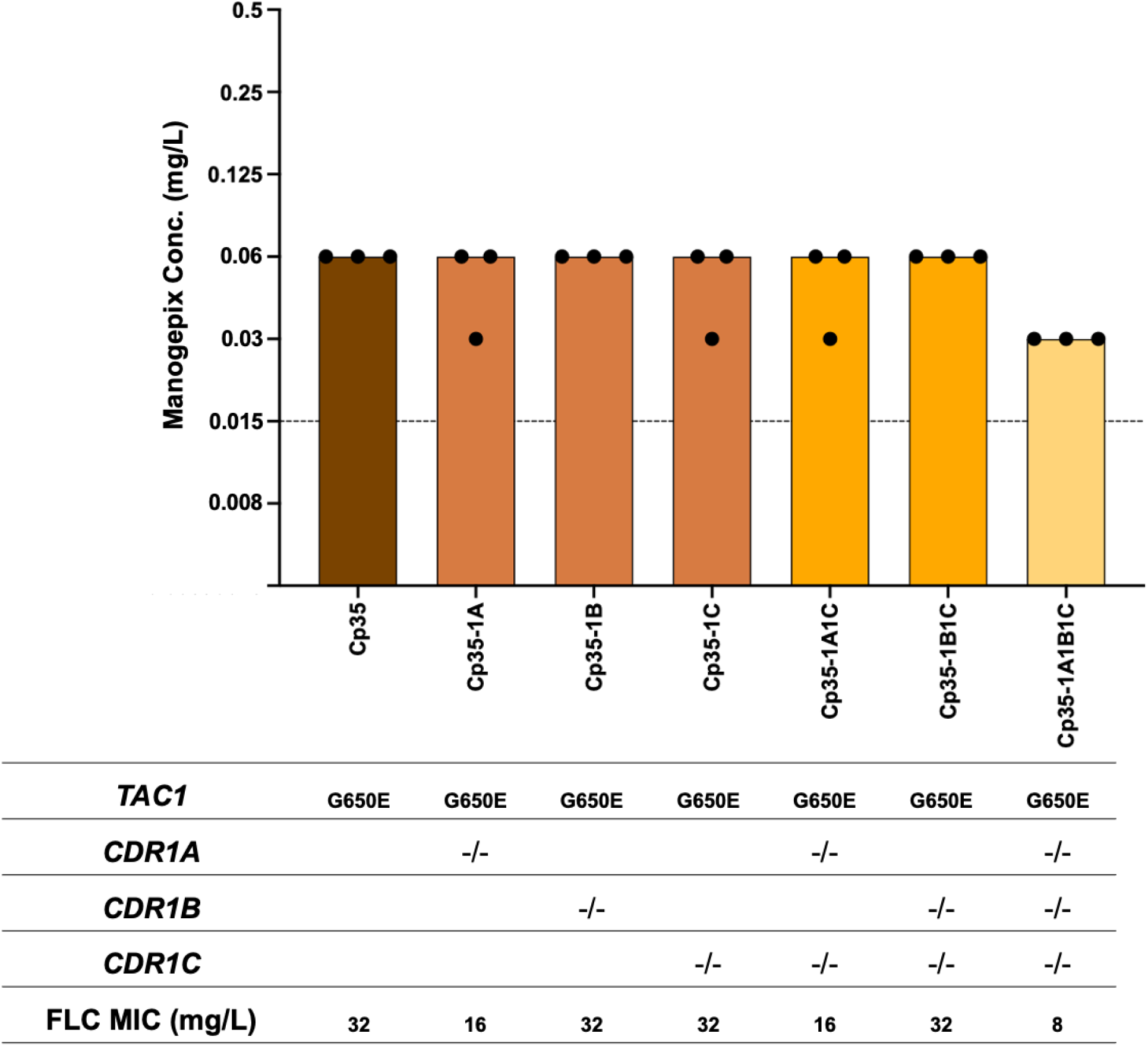
Manogepix MIC for (A) *C. parapsilosis* isolate Cp35 and its *TAC1*^WT^ derivative strain, and (B) Cp35 and its derivative strains in which *CDR1A*, *CDR1B*, *CDR1C* have been knocked out individually or in combination. Each bar in graph represents the modal MIC value, and each dot represents the measurements from three independent assays. Proposed epidemiologic cutoff value is indicated by the dashed line.

### Reduced susceptibility to manogepix in *C. parapsilosis* due to mutations in *TAC1* are driven in part by *CDR1A* and *CDR1C*

Fluconazole resistance in *C. parapsilosis* can be due in part to mutations in *TAC1* which drive overexpression of *CDR1A*, *CDR1B*, and *CDR1C*. We measured manogepix MICs in the fluconazole-resistant isolate Cp35, which carries a *TAC1*^G650E^ mutation, and its derivatives disrupted for *CDR1A*, *CDR1B*, or *CDR1C*, respectively (**Fig 4B**). Disruption of each of these ABC transporter genes alone, or the combination of *CDR1C* with either *CDR1A* or *CDR1B* had no effect on manogepix MIC. Further, the disruption of all three transporter genes decreased manogepix MIC by only one dilution, suggesting there are additional *TAC1* target genes contributing to *TAC1*-mediated reduced manogepix susceptibility.

### A mutation in *PDR1* that confers resistance to fluconazole in *C. glabrata* confers reduced susceptibility to manogepix

To determine if *PDR1* activating mutations that confer fluconazole resistance in *C. glabrata* influence susceptibility to manogepix, we first measured manogepix susceptibilities in susceptible isolate SM1, its matched fluconazole-resistant isolate SM3, which carries a *PDR1*^L946S^ mutation, and a derivative of SM1 engineered to carry the *PDR1* allele from isolate SM3 (**Fig 5A**). Both SM3 and the SM1 derivative carrying the *PDR1*^SM3^ allele exhibited a one-dilution increase in manogepix MIC to 0.06 µg/mL as compared to SM1 with an MIC of 0.03 µg/mL. These findings indicate that activating mutations in *PDR1* that confer fluconazole resistance in *C. glabrata* likewise result in reduced susceptibility to manogepix.

**Figure 5.**
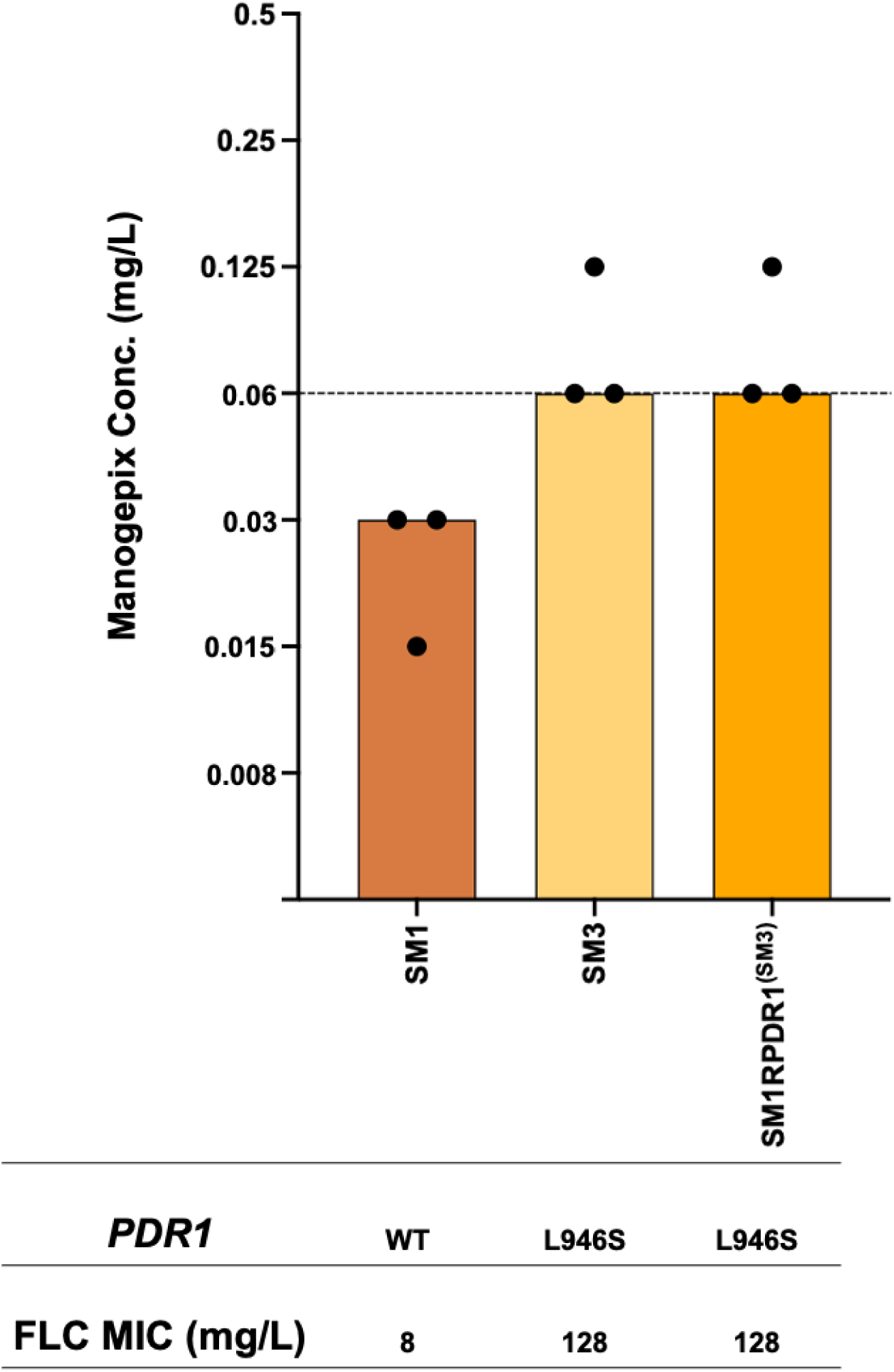
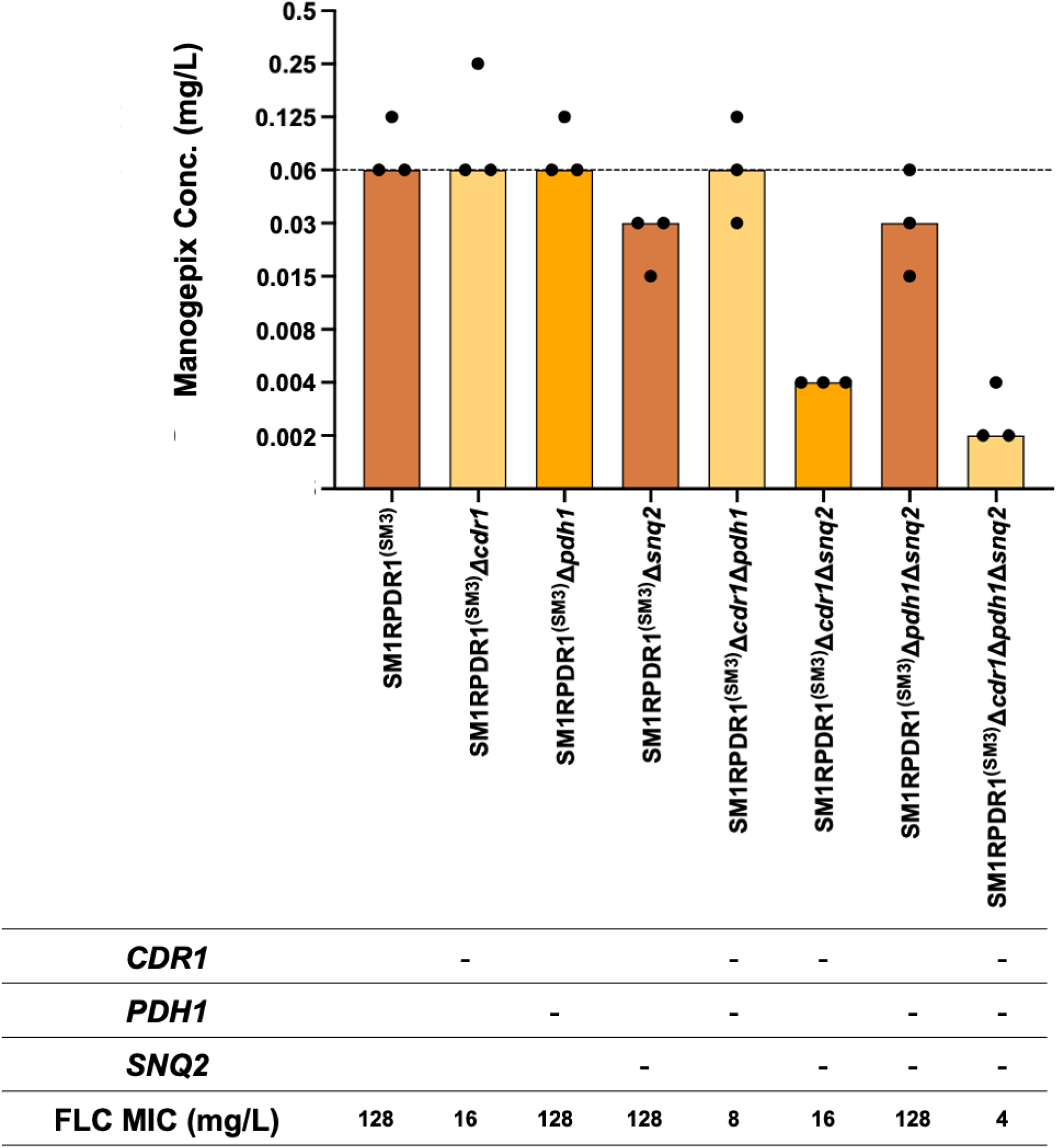
Manogepix MIC for (A) *C. glabrata* clinical isolate matched set SM1 and SM3 and SM1 derivative strain harboring *PDR1* allele from SM3, and (B) SM1 derivative strain harboring *PDR1* allele from SM3 and its derivative strains in which *CDR1*, *PDH1*, *SNQ2* have been knocked out individually or in combination. Each bar in graph represents the modal MIC value, and each dot represents the measurements from three independent assays. Proposed epidemiologic cutoff value is indicated by the dashed line.

### *SNQ2* and to a lesser extent *CDR1* and *PDH1* are drivers of reduced susceptibility to manogepix in *C. glabrata*

The *PDR1*^L946S^ mutation in isolate SM3 has been shown to drive fluconazole resistance through up-regulation of the genes encoding the Cdr1, Pdh1, and Snq2 ABC transporters (23). We therefore measured manogepix susceptibilities in strains derived from the *PDR1*^SM3^ derivative of isolate SM1 that lack *CDR1*, *PDH1*, and *SNQ2* (**Fig 5B**). Deletion of either *CDR1* or *PDH1* alone in this strain had no effect on manogepix MIC, whereas deletion of *SNQ2* increased susceptibility by one dilution from 0.06 µg/mL to 0.03 µg/mL. Deletion of both *CDR1* and *PDH1* had no effect (MIC 0.06 µg/mL), deletion of both *CDR1* and *SNQ2* together by 4 dilutions to 0.004 µg/mL, and deletion of all three transporters together by five dilutions to 0.002 µg/mL. These data indicate that *SNQ2* is the predominant driver of reduced susceptibility of manogepix due to this *PDR1* mutation.

## Discussion

Fosmanogepix is among several antifungal agents currently in late stages of development and has shown promise in a phase II study of non-neutropenic patients with candidemia (24). Promising results have also been reported in the treatment of a limited number of ICU patients with *C. auris* candidemia. While it has generally shown potent in vitro activity against *Candida* species, a correlation to fluconazole susceptibilities has been observed in some studies (8). Our findings suggest that this correlation might be explained by mutations in the transcription factors that drive overexpression of ABC transporters in these species.

A *TAC1B*^N865D^ mutation was recently implicated in decreased manogepix susceptibility in a laboratory-evolved clade I *C. auris* strain (17). This strain exhibited increased *CDR1* expression, and loss of either *CDR1* or *TAC1B* conferred increased susceptibility. In the present study, we observed a similar reduction in manogepix susceptibility in fluconazole-resistant isolates and strains harboring different *TAC1B* mutations. In addition to *CDR1*, these mutations have been shown to drive increased expression of *MDR1* which encodes a MFS transporter involved in fluconazole resistance. We found that changes in manogepix susceptibility due to these *TAC1B* mutations were due to overexpression of *CDR1*, not *MDR1*. Unlike mutations identified in fluconazole-resistant isolates, the *TAC1B*^N865D^ mutation generated in manogepix-evolved strains resulted in increased *CDR1* expression but not *MDR1* expression suggesting manogepix may preferentially select for *TAC1B* mutations that selectively regulate *CDR1* expression.

We also found that mutations in the zinc cluser transcription factor *TAC1* in *C. albicans* and *C. parapsilosis* that contribute to fluconazole resistance likewise result in decreased susceptibility to manogepix. In *C. albicans*, Tac1 has been shown to upregulate, via activating mutations or in response to inducers such as fluphenazine, the expression of the ABC transporter genes *CDR1* and *CDR2* and contribute to fluconazole resistance (10). It is likely that the reduced susceptibility to manogepix we observed in isolates and strains carrying *TAC1* mutations is due to overexpression of one or both of these transporters. However, one limitation of our study is the lack of *CDR1* or *CDR2* deletion mutants within our strain collection limiting our ability to test this hypothesis. The *TAC1*^G650E^ mutation in *C. parapsilosis* conferred reduced susceptibility, as disruption of these Cdr genes in the presence of this *TAC1* mutation conferred increased susceptibility. However, as disruption of *CDR1A*, *CDR1B*, and *CDR1C* together did not restore susceptibility to the level observed when the *TAC1*^G650E^ mutation was corrected to the wild-type sequence, it is likely that other Tac1-regulated genes contribute to this phenotype.

In *C. glabrata*, *PDR1* regulates the multidrug-resistant ABC transporter genes *CDR1*, *PDH1*, and *SNQ2*, all of which have been shown to contribute to fluconazole resistance. Importantly, fluconazole resistance in *C. glabrata* is almost exclusively due to such mutations. We found that the *PDR1*^L946S^ mutation which confers fluconazole resistance in a clinical *C. glabrata* isolate also confers reduced susceptibility to manogepix. This was dependent in large part on *SNQ2*, but loss of *SNQ2*, *CDR1*, and *PDH1* in combination conferred hypersusceptibility to manogepix, revealing a possible strategy for sensitizing *C. glabrata* to this antifiungal.

Our findings demonstrate that, like the *TAC1B*^N865D^ mutation leading to the substitution conferring reduced manogepix susceptibility in the *C. auris* manogepix-evolved laboratory strain, mutations in this and similar transcription factor genes that confer fluconazole resistance in clinical isolates of multiple *Candida* species likewise confer reduced manogepix susceptibility. This may explain previously observed correlations between fluconazole and manogepix susceptibilities and may have clinical implications for treating fluconazole-resistant isolates that harbor such mutations.

## Materials and Methods

### Strain and isolate growth conditions

The strains and isolates used in this study are listed in **Table 1** and were routinely propagated in YPD (1% yeast extract, 2% peptone, 2% dextrose) at 30°C for all species except for *C. auris* (35°C) and stored in 40% glycerol at -80°C.

**Table 1.**
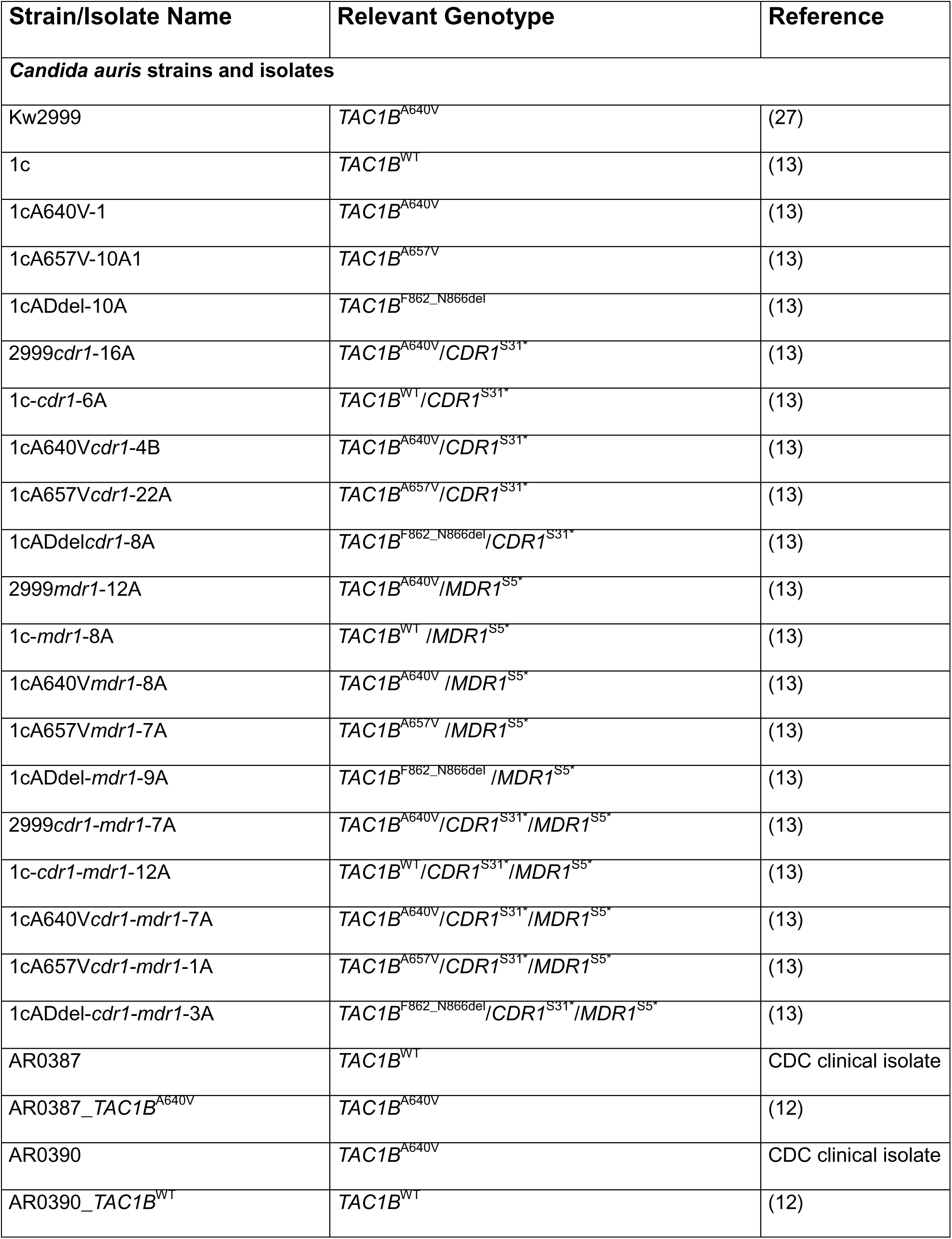
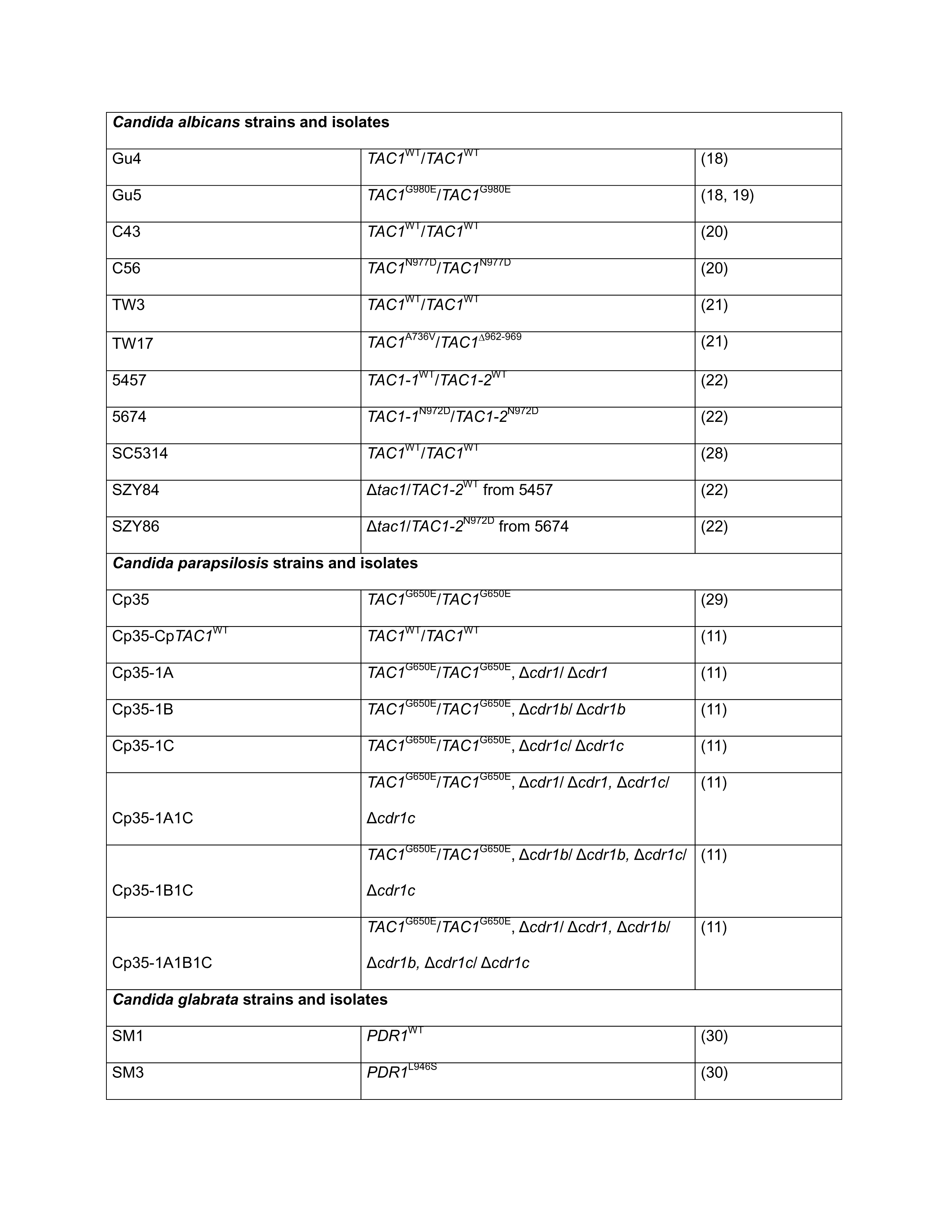
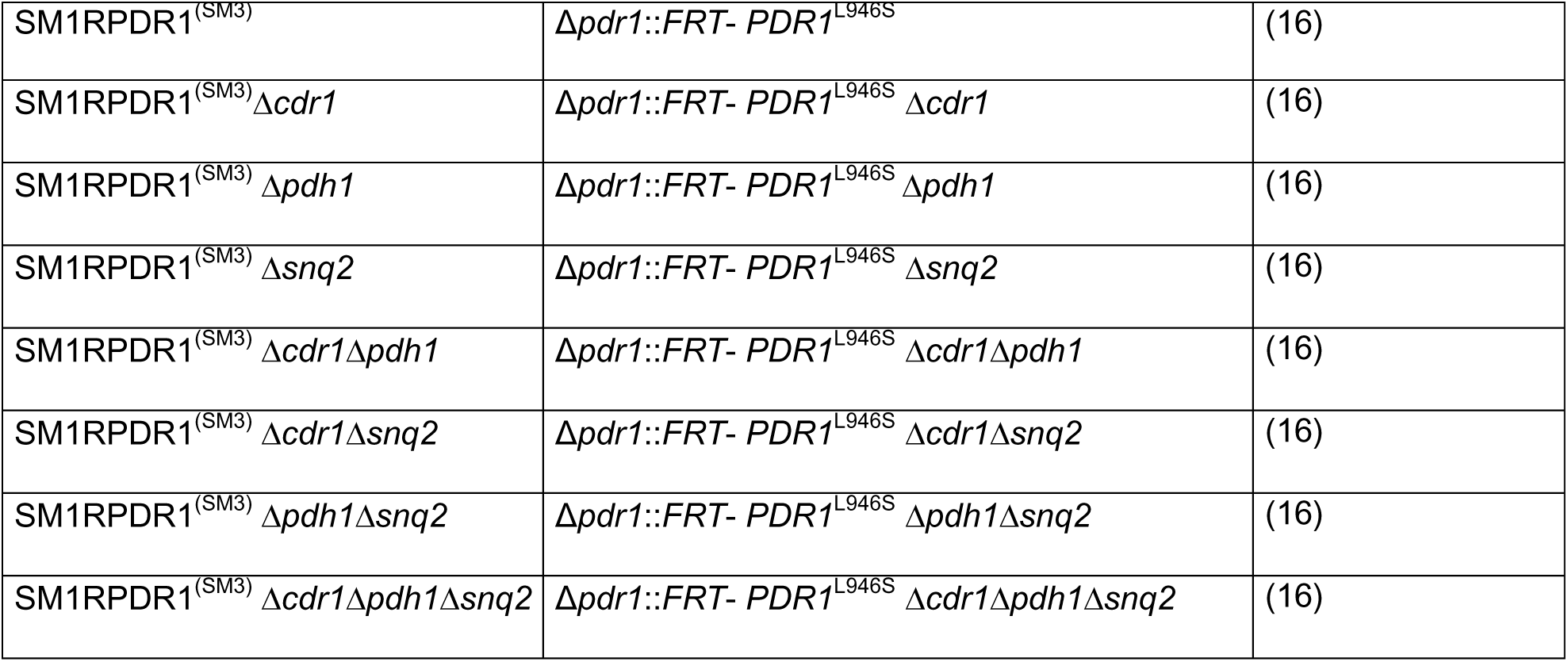

### Broth microdilution assays

Antifungal susceptibility testing was performed by broth microdilution according to the methods published in the CLSI M27 standard (25, 26). Manogepix stocks were prepared in DMSO with further dilutions in RPMI-1640 buffered with 0.165M MOPS (pH 7.0). The concentration range of manogepix that was tested was 0.002 to 1 μg/ml, and the final DMSO concentration per well was 1% v/v. Trays were incubated at 35°C for 24 hours, and the manogepix MIC was read as the lowest concentration that resulted in at least 50% inhibition of growth compared to the drug-free growth control well. Each isolate was tested on three separate days. *Candida parapsilosis* ATCC 22019 and *Candida albicans* ATCC 90028 served as the quality control isolates, as recommended by CLSI, and were included on each day of testing.

## Acknowledgements

This work was supported by NIH award R01 AI169066 to P.D.R. and C.A.C. and in part by the National Cancer Institute of the National Institutes of Health under Award Number P30 CA021765 awarded to the Hartwell Center at St. Jude Children’s Research Hospital. The funders had no role in study design, data collection and interpretation, or the decision to submit the work for publication.

